# Genomic Bayesian Prediction Model for Count Data with Genotype × Environment Interaction

**DOI:** 10.1101/034967

**Authors:** Abelardo Montesinos-López, Osval A. Montesinos-López, José Crossa, Juan Burgueño, Kent M. Eskridge, Esteban Falconi-Castillo, Xingyao He, Pawan Singh, Karen Cichy

## Abstract

Genomic tools allow the study of the whole genome and are facilitating the study of genotype-environment combinations and their relationship with the phenotype. However, most genomic prediction models developed so far are appropriate for Gaussian phenotypes. For this reason, appropriate genomic prediction models are needed for count data, since the conventional regression models used on count data with a large sample size (*n*) and a small number of parameters (*p*) cannot be used for genomic-enabled prediction where the number of parameters (*p*) is larger than the sample size (*n*). Here we propose a Bayesian mixed negative binomial (BMNB) genomic regression model for counts that takes into account genotype by environment (*G* × *E*) interaction. We also provide all the full conditional distributions to implement a Gibbs sampler. We evaluated the proposed model using a simulated data set and a real wheat data set from the International Maize and Wheat Improvement Center (CIMMYT) and collaborators. Results indicate that our BMNB model is a viable alternative for analyzing count data.

## Introduction

A phenotype is the result of genotype (*G*), environment (*E*) and the genotype by environment interactions (*G × E*) in most living organisms. Garrod (1902) observed that the effect of genes on phenotype could be modified by the environment (E). Similarly, Turesson (1922) demonstrated that the development of a plant is often influenced by its surroundings. He postulated the existence of a close relationship between crop plant varieties and their environment, and stressed that the presence of a particular variety in a given locality is not just a chance occurrence; rather, there is a genetic component that helps the individual adapt to that area.

For these reasons, today the consensus is that *G × E* is useful for understanding genetic heterogeneity under different environmental exposures (Kraft et al., 2007; Van Os and Rutten, 2009) and for identifying high-risk or productive subgroups in a population (Murcray et al., 2009); it also provides insight into the biological mechanisms of complex traits such as disease resistance and yield (Thomas, 2011), and improves the ability to discover resistance genes that interact with other factors that have little marginal effects (Thomas, 2011). However, finding significant *G × E* interactions is challenging. Model misspecification, inconsistent definition of environmental variables, and insufficient sample sizes are just a few of the issues that often lead to low-power and non-reproducible findings in *G × E* studies (Jiao et al., 2013; Winham and Biernacka, 2013).

Genomics and its breeding applications are developing very quickly with the goal of predicting yet-to-be observed phenotypes or unobserved genetic values for complex traits and inferring the underlying genetic architecture utilizing large collections of markers (Goddard and Hayes, 2009; Zhang et al., 2014). Also, genomics is useful when dealing with complex traits that are multi-genic in nature and have major environmental influence (Perez-de-Castro et al., 2012). For these reasons, the use of whole genome prediction models continues to increase. In genomic prediction, all marker effects are fitted simultaneously on a model and simulation studies promote the use of this methodology to increase genetic progress in less time. For continuous phenotypes, models have been developed to regress phenotypes on all available markers using a linear model (Goddard and Hayes, 2009; de los Campos et al., 2013). However, in plant breeding, the response variable in many traits is a count (y=0,1,2,…), for example, number of panicle per plant, number of seed per panicle, weed count per plot, etc. Count data are discrete, non-negative, integer-valued, and typically have right-skewed distributions (Yaacob et al., 2010).

Poisson regression and negative binomial regression are often used to deal with count data. These models have a number of advantages over an ordinary linear regression model, including a skewed, discrete distribution (0,1,2,3,…,) and the restriction of predicted values for phenotypes to non-negative numbers (Yaacob et al., 2010). These models are different from an ordinary linear regression model. First, they do not assume that counts follow a normal distribution. Second, rather than modeling *y* as a linear function of the regression coefficients, they model a function of the response mean as a linear function of the coefficients (Cameron and Trivedi, 1986). Regression models for counts are usually nonlinear and have to take into consideration the specific properties of counts, including discreteness and non-negativity, and are often characterized by overdispersion (variance greater than the mean) (Zhou et al., 2012).

However, in the context of genomic selection, it is still common practice to apply linear regression models to these data or to transformed data (Montesinos-López et al., 2015a,b). This does not take into account that: (a) many distributions of count data are positively skewed, many observations in the data set have a value of 0, and the high number of 0’s in the data set does not allow a skewed distribution to be transformed into a normal one (Yaacob et al., 2010); and (b) it is quite likely that the regression model will produce negative predicted values, which are theoretically impossible (Yaacob et al., 2010; Stroup, 2015). When transformation is used, it is not always possible to have normally distributed data and many times transformations not only do not help, they are counterproductive. There is also mounting evidence that transformations do more harm than good for the models required by the vast majority of contemporary plant and soil science researchers (Stroup, 2015). To the best of our knowledge, only the paper of Montesinos-López et al. (2015c) is appropriate for genomic prediction for count data under a Bayesian framework; however it does not take into account *G × E* interaction.

In this paper, we extend the NB regression model for counts proposed by Montesinos-López et al. (2015c) to take into account *G × E* by using a data augmentation approach. A Gibbs sampler was derived since all full conditional distributions were obtained, which allows drawing samples from them to estimate the required parameters. In addition, we provide all the details of the efficient derived Gibbs sampler so it can be easily implemented by most plant and animal scientists. We illustrate our proposed methods with a simulated data set and a real data set on wheat Fusarium head blight. We compare our proposed models (NB and Poisson) with the Normal and Log-Normal models that are commonly implemented for analyzing count data. We also provide R code for implementing the proposed models.

## Materials and Methods

The data used in this study were taken from a Ph.D. thesis (Falconi-Castillo, 2014) aimed at identifying sources of resistance to Fusarium head blight (FHB), caused by *Fusarium graminearum* and identify genomic regions and molecular markers linked to FHB resistance through association analysis.

## Experimental data

### Phenotypic data

A total of 297 spring wheat lines developed by the International Maize and Wheat Improvement Center (CIMMYT) was assembled and evaluated for resistance to *F. graminearum* in México over two years (2012 and 2014) and Ecuador for one year (2014). In this paper we used only 182 spring wheat lines since only for these lines we have complete marker information.

### Genotypic data

DNA samples were genotyped using an Illumina 9K SNP chip with 8,632 SNPs (Cavanagh et al., 2013). SNP markers with unexpected genotype AB (heterozygous) were recoded as either AA or BB based on the graphical interface visualization tool of the software GenomeStudio^®^ (Illumina). SNP markers that did not show clear clustering patterns were excluded. In addition, 66 simple sequence repeats (SSR) markers were screened. After filtering the markers for the minor allele frequency (MAF) of 0.05 and deleting markers with more than 10% of no calls, the final set of SNPs was of 1,635 SNP.

### Data and software availability

The phenotypic (FHB) and genotypic (marker) data used in this study as well as basic R codes (R Core Team, 2015) for fitting the models can be directly downloaded from the repository at http://hdl.handle.net/11529/10575

## Statistical Models

We used *y_ijt_* to represent the count response for the tth replication of the *j*th line in the *i*th environment with *i* = 1, *…*,*I*; *j* = 1,2, *…*,*J*,*t* = 1,2, *…*,*n_ij_* and we propose the following linear predictor that takes into account *G × E:*

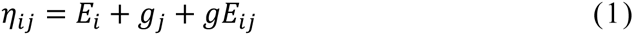

where *E_i_* represents the environment *i*, *g_j_* is the marker effect of genotype *j*, and *gE_ij_* is the interaction between markers and environment; *I* = 3, since we have three environments (Batan 2012, Batan 2014, and Chunchi 2014), *J* = 182, since it is the number of lines under study, and *n_ij_* represents the number of replicates of each line in each environment (the minimum and maximum *n_ij_* found per line were 10 and 20). The number of observations in each environment *i* is 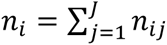, while the total number of observations is 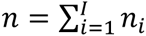. is the product of the number of environments and number of lines. Four models were implemented using the linear predictor given in expression (1).

### Model NB

Distributions: 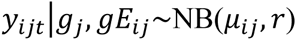, with *r* being the scale parameter, *μ_ij_* = exp(*ηij*), 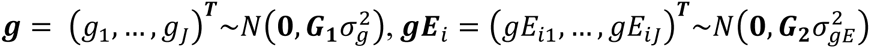. Note that the NB distribution has expected value *μ_ij_* and is smaller than the variance 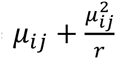. ***G*_1_** and ***G*_2_** were assumed known, with ***G*_1_** computed from marker ***X*** *data* (for k = 1,…, p markers) as 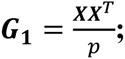 this matrix is called the Genomic Relationship Matrix (GRM) (VanRaden, 2008). While ***G***_2_ is computed as ***G*_2_** = ***I_I_*** ⊗ ***G***_1_ of order *IJ*x*IJ* and ⊗ denotes the Kronecker product, *I_I_* means that we assume independence between environments.

### Model Pois

This model is the same as **Model NB**, except that *y_ijt_/g_j_*, *gE_ij_~Poisson*(*μ_ij_*). Since according to Zhou et al. (2012) and Teerapabolarn and Jaioun (2014) the 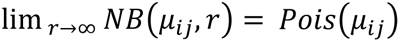, **Model Pois** was implemented using the same method as **Model NB**, but fixing *r* to a large value, depending on the mean count. We used *r* = 1000, which is a good choice when the mean count is less than 100.

### Model Normal

Model Normal is similar to **Model NB**, except that 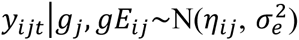 with identity link function.

### Model Log-Normal

Model Log-Normal is similar to **Model NB**, except that 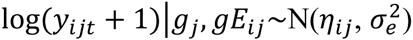 with identity link function.

When *p > n*, implementing **Models NB** and **Pois** is challenging. For this reason, we propose a Bayesian method for dealing with situations when *p > n*. The **Models Normal** and **Log-Normal** were implemented in the package BGLR of de los campos et al. (2014).

#### Bayesian mixed negative binomial regression

Rewriting the linear predictor (1) as 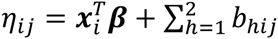, with 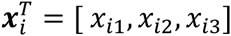, where *x_ik_* is an indicator variable that takes the value of 1 if it is observed in environment *i* and 0 otherwise, for *k* = 1,2,3; ***β****^T^* = [*β*_1_,*β*_2_,*β*_3_,] since three is the number of environments under study, *b*_1_*_ij_ = g_j_* and *b*_2_*_ij_ = gE_ij_*. Note that in a sequence of independent Bernoulli (*π_ij_*) trials, the random variable *y_ijt_* denotes the number of successes before the r*th* failure occurs. Then

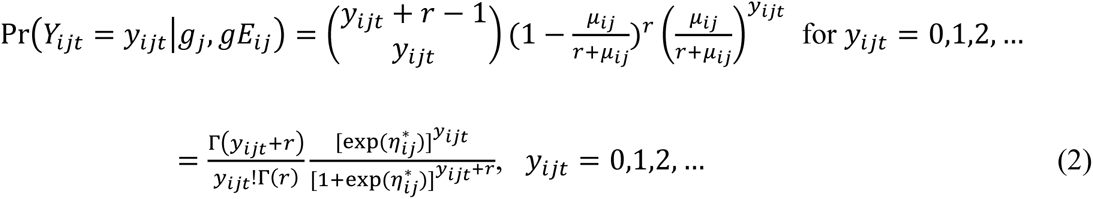

Since 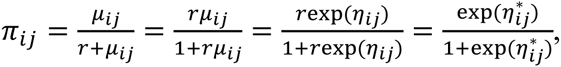 where 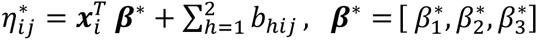 with 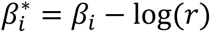 since 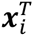 is composed of three indicator variables. We can rewrite (Eq 2) as:

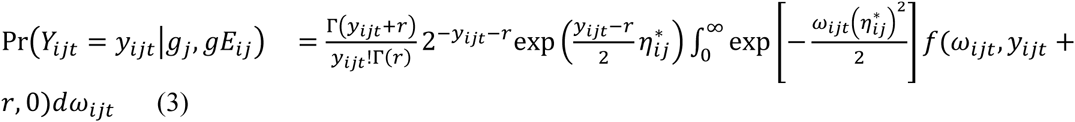

Expression (3) was obtained using the equality given by Polson et al. (2013): 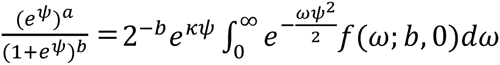, where *κ = a − b/*2 and *f*(.,*b*, 0) denotes the density of *PG*(*b*, *c* = 0), the *PG* Pólya-Gamma distribution with parameters *b* and *c* = 0 (see Definition 1 in Polson et al., 2013).

From here, conditional on 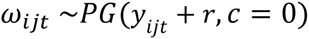,

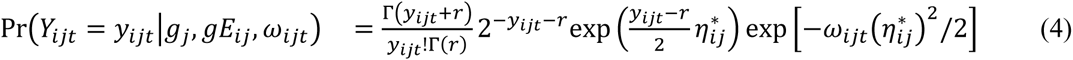

To be able to get the full conditional distributions, we provide the prior distributions, *f*(***θ***), for all the unknown model parameters 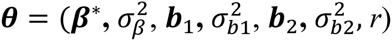. We assume prior independence between the parameters, that is,

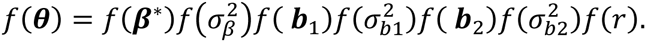

We assign conditionally conjugate but weakly informative prior distributions to the parameters because we have no prior information. Prior specification in terms of ***β****^*^* instead of ***β*** is for convenience. We adopt proper priors with known hyper-parameters whose values we specify in model implementation to guarantee proper posteriors. We assume that 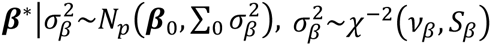 where 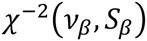 denotes a scaled inverse chi-square distribution with shape *v_β_* and scale *S_β_* parameters, 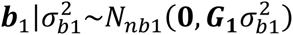, 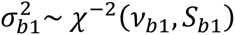, 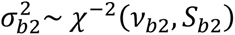 and 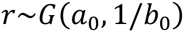. Next we combine (Eq 4) using all data with priors to get the full conditional distribution for parameters ***β****^*^*, 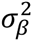, ***b***_1_, 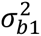, ***b***_2_, 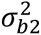 and *r*.

#### Full conditional distributions

The full conditional distribution of ***β****^*^* is given as:

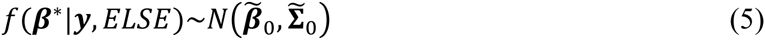

where 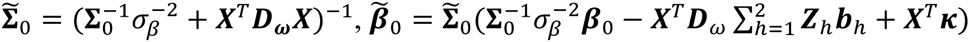, 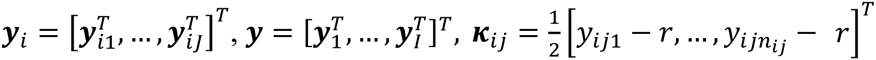, 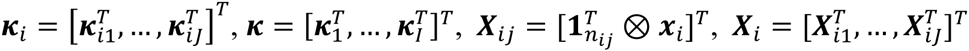, 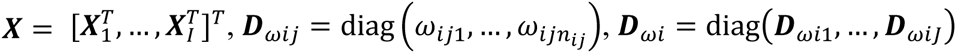, 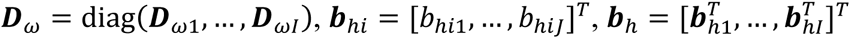, 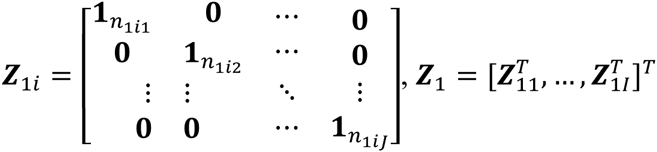, and ***Z***_2_ = ***Z***_1_ * ~ ***X*** where * ~ indicates the horizontal Kronecker product between ***Z***_1_ and ***X***. The horizontal Kronecker product performs a Kronecker product of ***Z***_1_ and ***X*** and creates a new matrix by stacking these row vectors into a matrix. ***Z***_1_ and ***X*** must have the same number of rows, which is also the same number of rows in the result matrix. The number of columns in the result matrix is equal to the product of the number of columns in ***Z***_1_ and ***X***. When the prior for ***β****^*^* **∝** constant, the posterior distribution of ***β****^*^* is also normally distributed, 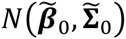, but we set the term 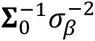 to zero in both 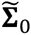 and 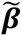.

The fully conditional distribution of *ω_ijt_* is

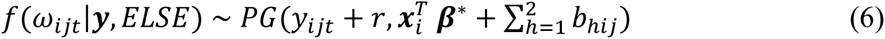

Defining ***η****^h^* = ***X β****^*^* + ***Z****_h_****b****_h_*, with *h* = 1,2, the conditional distribution of ***b****_h_* is given as

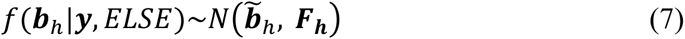

If ***η***^1^ = ***X β****^*^* + ***Z***_2_***b***_2_, then 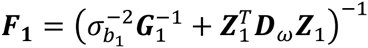, 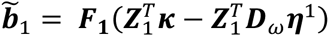 and then 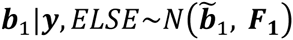. Similarly, by defining ***η***^2^ = ***X β****^*^* + ***Z***_1_***b***_1_, we arrive at the full conditional of ***b***_2_ as ***b***_2_|***y***, 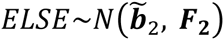, where 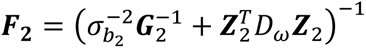, 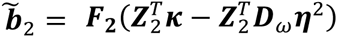.

The fully conditional distribution of 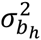, for *h* = 1,2, is

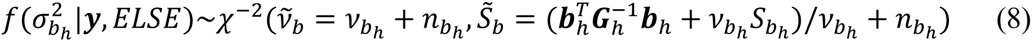

with 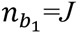 and 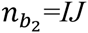.

The conditional distribution of 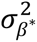 is

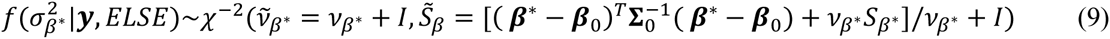

Taking advantage of the fact that the NB distribution can also be generated using a Poisson representation (Quenouille, 1949) as 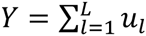 where 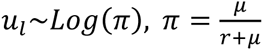 and is independent of *L~ Pois*(*−r*log(1 *−π*)), where *Log* and *Pois* denote logarithmic and Poisson distributions, respectively. Then we infer a latent count *L* for each *Y ~ NB*(*μ*, *r*) conditional on *Y* and *r*. Therefore, following Zhou et al. (2012), we obtain the full conditional of *r* by alternating

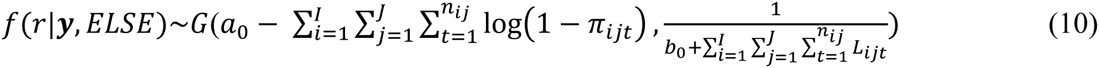

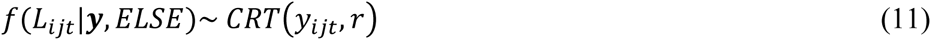

where *CRT*(*y_ijt_*,*r*) denotes a Chinese restaurant table (CRT) count random variable that can be generated as 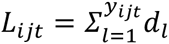, where 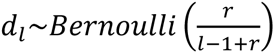. For details of the CRT random variable derivation, see Zhou and Carin (2012, 2015).

### Gibbs sampler

The Gibbs sampler for the latent parameters of the NB with *G × E* can be implemented by sampling repeatedly from the following loop:

1. Sample *ω_ijt_* values from the Pólya-Gamma distribution in (6).
2. Sample *L_ijt_~CRT*(*y_ijt_*,*r*) from (11).
3. Sample the scale parameter (*r*) from the gamma distribution in (10).
4. Sample the location effects (***β****^*^*) from the normal distribution in (5).
5. Sample the random effects (***b****_h_*) with *h* = 1,2, from the normal distribution in (7).
6. Sample the variance effects 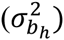 with *h* = 1,2, from the scaled inverted *χ*^2^ distribution in (8).
7. Sample the variance effect 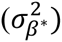 from the scaled inverted *χ*^2^ distribution in (9).
8. Return to step 1 or terminate when chain length is adequate to meet convergence diagnostics.

### Model implementation

The Gibbs sampler described above for the BMNB model was implemented in R-Core Team (2015). Implementation was done under a Bayesian approach using Markov Chain Monte Carlo (MCMC) through the Gibbs sampler algorithm, which samples sequentially from the full conditional distribution until it reaches a stationary process, converging with the joint posterior distribution (Gelfand and Smith, 1990). To decrease the potential impact of MCMC errors on prediction accuracy, we performed a total of 60,000 iterations with a burn-in of 30,000, so that 30,000 samples were used for inference. We did not apply thinning of the chains following the suggestions of Geyer (1992), MacEachern and Berliner (1994) and Link and Eaton (2012), who provide justification of the ban on subsampling MCMC output for approximating simple features of the target distribution (e.g., means, variances, and percentiles). We implemented the prior specification given in the section Bayesian mixed negative binomial regression with 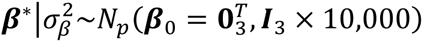, 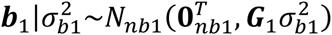, where ***G***_1_ is the GRM, that is, the covariance matrix of the random effects, 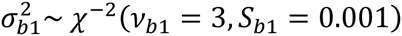, 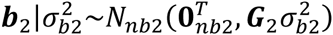, ***G***_2_ is the covariance matrix of the random effects that belong to the *G × E* term, 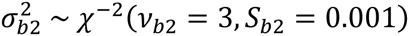, and *r~G*(*a*_0_ = 0.01,1*/*(*b*_0_ = 0.01)). All these hyper-parameters were chosen to lead weakly informative priors. The convergence of the MCMC chains was monitored using trace plots and autocorrelation functions. We also conducted a sensitivity analysis on the use of the inverse gamma priors for the variance components and we observed that the results are robust under different choices of priors.

### Assessing prediction accuracy

We used cross-validation to compare the prediction accuracy of the proposed models for count phenotypes. We implemented a 10-fold cross validation, that is, the data set was divided into 10 mutually exclusive subsets; each time we used 9 subsets for the training set and the remaining one for validation set. The training set was used to fit the model and the validation set was used to evaluate the prediction accuracy of the proposed models. To compare the prediction accuracy of the proposed models, we calculated the Spearman correlation (Cor) and the mean square error of prediction (MSEP), both calculated using the observed and predicted response variables of the validation set. Models with large absolute values of Cor indicate better prediction accuracy, while small MSEP indicate better prediction performance. The predicted observations, 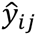, were calculated with *M* collected Gibbs samples after discarding those of the burn-in period. For **Models NB** and **Pois** the predicted values were calculated as 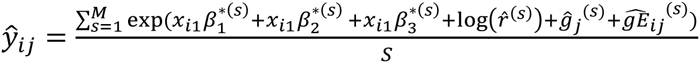, where 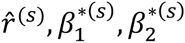, 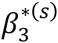 and 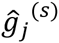 and 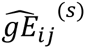 are estimates of 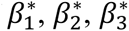, *r*, *g_j_* and *gE_ij_*, for line j in environment i obtained in the s*th* collected sample. For **Model Normal** as 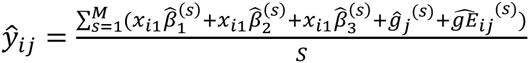 and for **Model LN** the predicted observations were calculated as 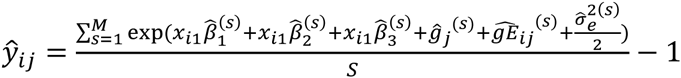, using the corresponding estimates of each model.

### Simulation study

To show the performance of the proposed Gibbs sampler for count phenotypes that takes into account *G × E*, we performed a simulation study under model (1) in two scenarios (S1 and S2). Scenario 1 had three environments (*I* = 3), 20 genotypes (*J* = 20), ***G***_1_ = ***I***_60_, ***G***_2_ = ***I_I_*** ⊗ ***G***_1_ and 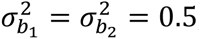, with four different numbers of replicates of each genotype in each environment, *n_ij_* = 5,10,20 and 40. Scenario 2 is equal to scenario 1, except that ***G***_2_ = 0.7***I***_60_ *+* 0.3***J***_60_, where ***J***_60_ is a square matrix of ones of order 60 *×* 60. In this second scenario, we imitated the correlation between lines of real data available in genomic selection. The priors used for the simulation study in both scenarios (S1 and S2) were approximately flat for all parameters: for 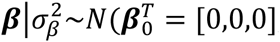, ***I***_3_ *×* 10000), for *r*~*G* (0.001,1/0.001), for 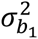 and 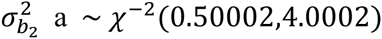, while for 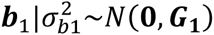, and for 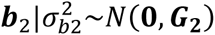. We computed 20,000 MCMC samples; Bayes estimates were computed with 10,000 samples since the first 10,000 were discarded as burning. We report average estimates obtained by using the proposed Gibbs sampler along with standard deviations (SD) (Table 1). All the results in Table 1 are based on 50 replications.

**Table 1.**
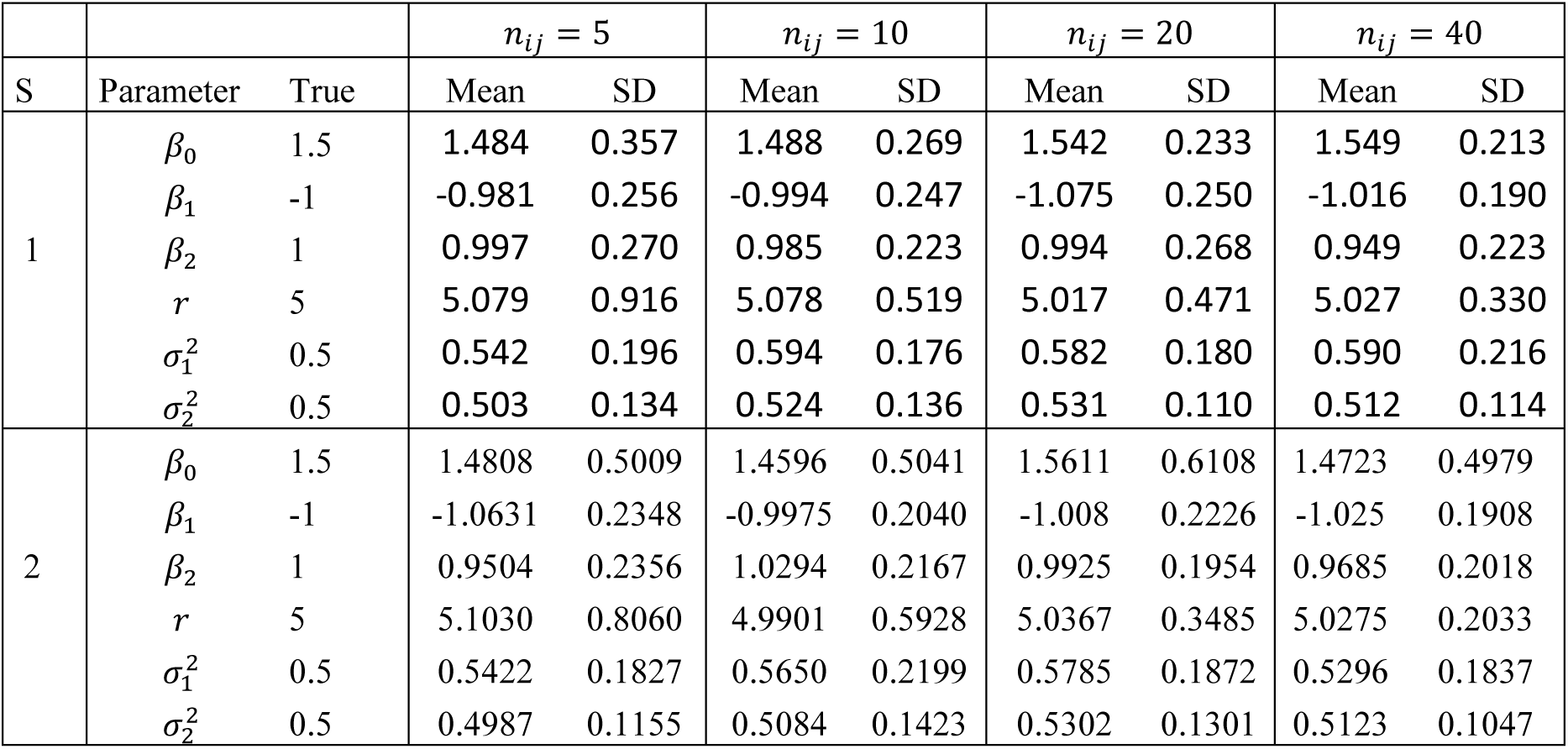
Posterior mean (Mean) and posterior standard deviation (SD) of the Bayesian method with four sample sizes (*n_ij_*) for Model NB. S denotes scenario.

## Results

Given in Table 1 are the results of the simulation study in both scenarios (S1 and S2). The bias when estimating the parameters is a little larger in S1 compared to S2. Also, parameter *β*_0_ is the parameter with larger bias (underestimated). Both variances 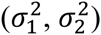 are overestimated in scenario 1, but only 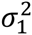 is overestimated in scenario 2. Also, with a sample size of *n_ij_* = 5, parameter *r* had a larger SD; however, for larger sample sizes (*n_ij_* = 20,40), the SD were considerably reduced. In general, there was not a large reduction in SD when the sample size increased from 5 to 10, 20 and 40, the exception being the estimation of *r* in both scenarios and the estimation of *β*_0_ in scenario 1, where there was a large reduction in SD when the sample size increased. Although estimations do not totally agree with the true values of the parameters, the proposed Gibbs sampler for count data that takes into account *G × E* did a good job of estimating the parameters, since the estimates are close to the true values with a SD of reasonable size.

Using the real data set, we compared four scenarios (given in Table 2) for each model. Table 2 shows that in the linear predictor, scenarios 1 and 2 do not take into account interaction effects, only main effects. Also, scenarios 1 and 3 do not use marker information. These four scenarios were studied to investigate the gain in model fit and prediction ability taking into account the interaction effects and using the marker information available.

**Table 2.**
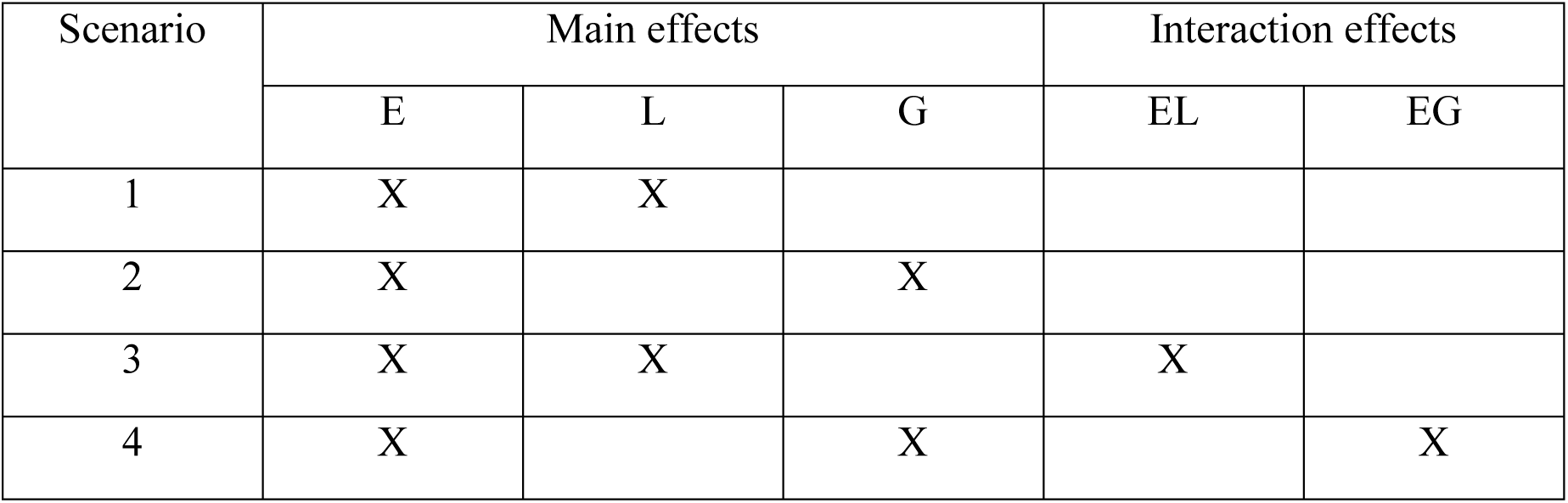
Scenarios proposed to fit the real data set with **Models NB**, **Pois, Normal and LN**. E stands for Environment, L for lines, G for lines taking into account markers; EL and EG are interaction effects of E and L and E and G.

The posterior means (Mean), posterior standard deviation (SD) of the scalar parameters, and posterior predictive checks for each scenario of the proposed models are given in Table 3. For the four models, the posterior means of the beta regression coefficients, variance components, and over-dispersion parameters (*r*) are similar between scenarios 1 and 2 and between scenarios 3 and 4. In terms of goodness of fit measured by the loglikelihood posterior mean (loglink), the scenarios rank as follows: scenario 3, rank 1; scenario 4, rank 2; scenario 1, rank 3; and scenario 2, rank 4, for the four proposed models, with the exception of **Model Pois** where the ranking was scenario 3, rank 1; scenario 4, rank 2; scenario 2, rank 3; and scenario 1, rank 4. Therefore, there is evidence that with the four proposed models in terms of goodness of fit, the best scenario is S3. Of the four models under study, Table 3 shows that **Model LN** reports the best fit since it has the largest Loglik.

**Table 3.**
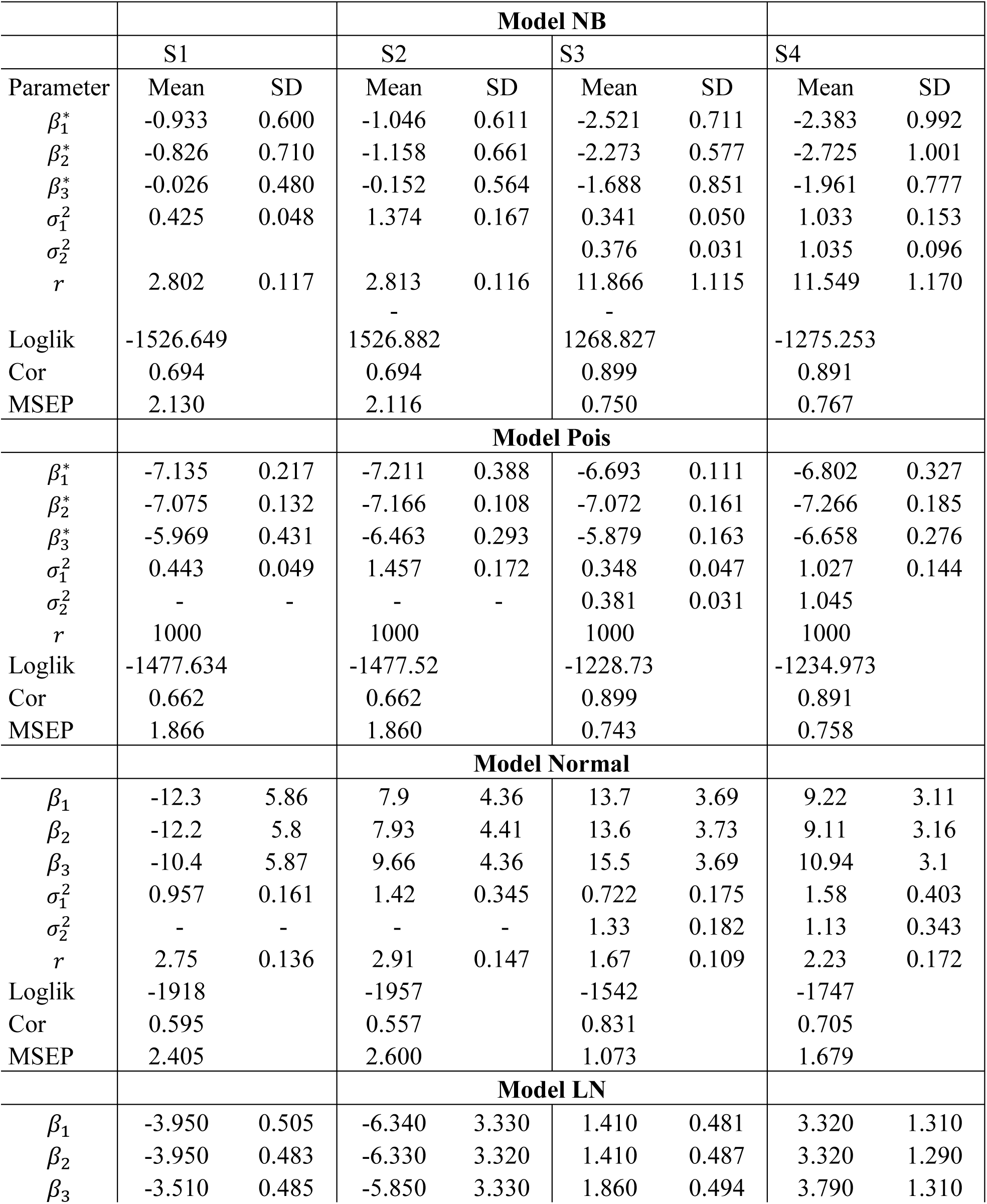
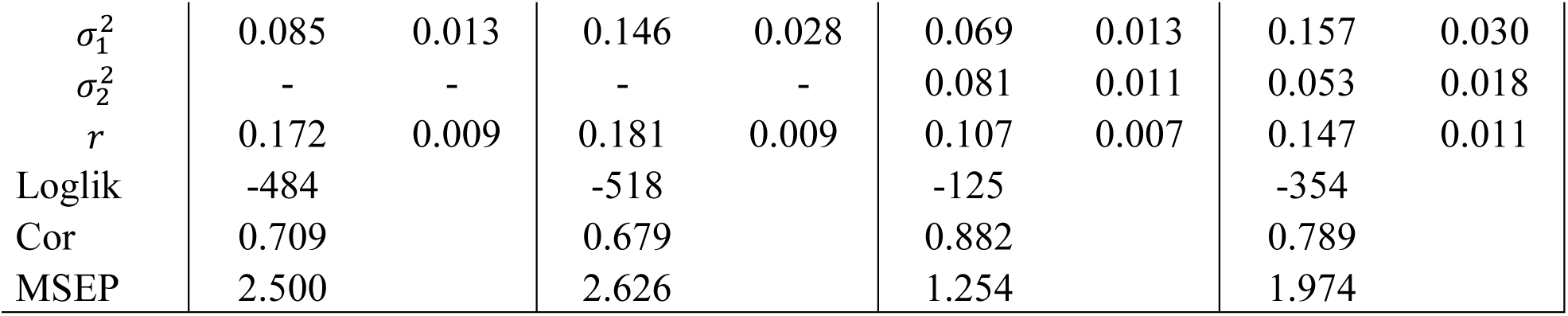
Estimated beta coefficients, variance components, and posterior predictive checks for the four scenarios (S1, S2, S3, S4) for each proposed model (**Model NB**, **Model Pois**, **Model Normal and Model LN**). Mean stands for posterior mean and SD for posterior standard deviation.

In Table 4 we present the mean and standard deviation of the posterior predictive checks (Cor and MSEP) for each location (Batan 2012, Batan 2014 and Chunchi 2014) resulting from the 10-fold cross-validation implemented for the four models and four scenarios. The predictive checks given in Table 4 were calculated using the testing set. In **Model NB**, according to the Spearman Correlation, the ranking of scenarios was as follows: in Batan 2012 and Batan 2014, 1 for scenario 4, 2 for scenario 3, 3 for scenario 1, and 4 for scenario 2. In Chunchi 2014, the ranking was 1 for scenario 3, 2 for scenario 2, 3 for scenario 4, and 4 for scenario 4. With the MSEP, the ranking for **Model NB** in Batan 2012 was 1 for scenario 3, 2 for scenario 4, 3 for scenario 1, and 4 for scenario 2. In Batan 2014, the ranking was 1 for scenario 2, 2 for scenario 1, 3 for scenario 3, and 4 for scenario 4. In Chunchi 2014, the ranking in terms of MSEP was 1 for scenario 3, 2 for scenario 2, 3 for scenario 4, and 4 for scenario 1. Under **Model Pois**, the ranking of the 4 scenarios in each locality was exactly the same as the ranking reported for **Model NB**. For **Model Normal** in terms of the Spearman correlation, scenario 1 was the best in prediction accuracy in Batan 2012 and Chunchi 2014, while scenario 4 was the worst in all three locations. In terms of MSEP, the best scenario was 3 in Batan 2014 and Chunchi 2014, and the worst was scenario 4 in Batan 2014 and Chunchi 2014. For **Model LN** in terms of the Spearman correlation, the best scenarios were scenarios 1 and 2, and the worst was scenario 3 in Batan 2012. In Batan 2014, the best scenario was 1, then scenario 3 and the worst was scenario 4. In Chunchi 2014, the best scenario was scenario 3, then scenario 2 and the worst was scenario 2. In terms of MSEP for Batan 2012, the best scenario was 3, then scenario 1 and the worst was scenario 4. In Batan 2014, the best scenario was 1, then 2 and the worst was scenario 4. Finally, in Chunchi 2014, the best scenario was 3, then 2 and the worst was scenario 1.

**Table 4.** Estimated posterior predictive checks with cross validation for **Models NB**, **Pois**, **Normal and LN**. () denotes the ranking of the four scenarios for each posterior predictive check. Each average was obtained as the mean of the rankings of the four posterior predictive checks for each scenario.

|  |  | Batan 2012 |  | Batan 2014 |  | Chunchi 2014 |  |
| --- | --- | --- | --- | --- | --- | --- | --- |
|  |  | Model NB |  |  |  |  |  |
| Scenario |  | Cor | MSEP | Cor | MSEP | Cor | MSEP |
| S1 | Mean | 0.426 (3) | 0.977 (3) | 0.427 (4) | 1.388 (2) | 0.182 (3) | 11.733 (4) |
|  | SD | 0.331 | 0.723 | 0.327 | 1.351 | 0.401 | 9.471 |
| S2 | Mean | 0.423 (4) | 0.980 (4) | 0.432 (3) | 1.383 (1) | 0.204 (2) | 11.222 (2) |
|  | SD | 0.327 | 0.717 | 0.325 | 1.356 | 0.373 | 8.614 |
| S3 | Mean | 0.539 (2) | 0.497 (1) | 0.522 (2) | 1.480 (3) | 0.224 (1) | 8.645 (1) |
|  | SD | 0.283 | 0.376 | 0.292 | 2.318 | 0.386 | 5.688 |
| S4 | Mean | 0.557 (1) | 0.607 (2) | 0.564 (1) | 1.850 (4) | 0.122 (4) | 11.343 (3) |
|  | SD | 0.243 | 0.438 | 0.222 | 2.684 | 0.407 | 8.154 |
|  |  | Model Pois |  |  |  |  |  |
| S1 | Mean | 0.426 (3) | 0.977 (3) | 0.427 (4) | 1.388 (2) | 0.182 (3) | 11.733 (4) |
|  | SD | 0.331 | 0.723 | 0.327 | 1.351 | 0.401 | 9.471 |
| S2 | Mean | 0.423 (4) | 0.980 (4) | 0.432 (3) | 1.383 (1) | 0.204 (2) | 11.222 (2) |
|  | SD | 0.327 | 0.717 | 0.325 | 1.356 | 0.373 | 8.614 |
| S3 | Mean | 0.539 (2) | 0.497 (1) | 0.522 (2) | 1.480 (3) | 0.224 (1) | 8.645 (1) |
|  | SD | 0.283 | 0.376 | 0.292 | 2.318 | 0.386 | 5.688 |
| S4 | Mean | 0.557 (1) | 0.607 (2) | 0.564 (1) | 1.850 (4) | 0.122 (4) | 11.343 (3) |
|  | SD | 0.243 | 0.438 | 0.222 | 2.684 | 0.407 | 8.154 |
|  |  | Model Normal |  |  |  |  |  |
| S1 | Mean | 0.358 (1) | 1.096 (4) | 0.367 (2) | 1.788 (1) | 0.148 (1) | 7.425 (2) |
|  | SD | 0.280 | 0.883 | 0.397 | 1.701 | 0.318 | 4.151 |
| S2 | Mean | 0.344 (2) | 0.988 (2) | 0.334 (3) | 2.010 (3) | 0.074 (3) | 7.454 (3) |
|  | SD | 0.326 | 0.652 | 0.440 | 2.462 | 0.330 | 4.339 |
| S3 | Mean | 0.330 (3) | 0.806 (1) | 0.371 (1) | 1.963 (2) | 0.146 (2) | 7.318 (1) |
|  | SD | 0.300 | 0.495 | 0.400 | 2.986 | 0.287 | 4.159 |
| S4 | Mean | 0.267 (4) | 1.029 (3) | 0.237 (4) | 2.373 (4) | 0.039 (4) | 8.482 (4) |
|  | SD | 0.338 | 0.731 | 0.445 | 3.420 | 0.238 | 4.326 |
|  |  | Model LN |  |  |  |  |  |
| S1 | Mean | 0.510 (1.5) | 0.661 (2) | 0.455 (1) | 1.601 (1) | 0.149 (2) | 8.099 (4) |
|  | SD | 0.208 | 0.419 | 0.307 | 2.348 | 0.379 | 5.113 |
| S2 | Mean | 0.510 (1.5) | 0.663 (3) | 0.433 (3) | 1.778 (2) | 0.086 (4) | 7.819 (2) |
|  | SD | 0.224 | 0.392 | 0.353 | 2.820 | 0.459 | 5.311 |
| S3 | Mean | 0.505 (3) | 0.639 (1) | 0.449 (2) | 1.871 (3) | 0.153 (1) | 7.759 (1) |
|  | SD | 0.208 | 0.451 | 0.313 | 3.162 | 0.371 | 5.209 |
| S4 | Mean | 0.428 (4) | 0.721 (4) | 0.427 (4) | 1.951 (4) | 0.087 (3) | 8.038 (3) |
|  | SD | 0.246 | 0.415 | 0.327 | 3.148 | 0.413 | 5.187 |

Table 5 gives the average of the ranks of the two posterior predictive checks (Cor and MSEP) that were used. Since we are comparing four scenarios for each model, the values of the ranks range from 1 to 4, and the lower the values, the better the scenario. For ties we assigned the average of the ranges that would have been assigned had there been no ties. Table 5 shows that the best scenarios were scenarios 3 and 4 under **Model NB** and **Pois** in Batan 2012. In Batan 2014 under **Models NB** and **Pois**, the best scenario was 3, while in Chunchi 2014, the best scenarios were 3 and 1. Under **Model Normal**, the best scenario was scenario 3 in Batan 2014 and Chuchi 2014, while in Batan 2012, the best scenarios were 2 and 3. Finally, under **Model LN**, the best scenario was 3 in Chunchi 2014, and scenario 1 in Batan 2012 and Batan 2014.

**Table 5.** Rank averages for the four scenarios for each model (**Models NB**, **Pois, Normal and LN**) resulting from the 10-fold cross-validation implemented. Each average was obtained as the mean of the rankings given in Table 4 for the two posterior predictive checks (Cor and MSEP) in each scenario.

| Scenario | Batan 2012 | Batan 2014 | Chunchi 2014 | Batan 2012 | Batan 2014 | Chunchi 2014 |
| --- | --- | --- | --- | --- | --- | --- |
|  | **Model NB** |  |  | **Model Normal** |  |  |
| S1 | 3 | 3 | 3.5 | 2.5 | 1.5 | 1.5 |
| S2 | 4 | 2 | 2 | 2 | 3 | 3 |
| S3 | 1.5 | 2.5 | 1 | 2 | 1.5 | 1.5 |
| S4 | 1.5 | 2.5 | 3.5 | 3.5 | 4 | 4 |
|  | **Model Pois** |  |  | **Model LN** |  |  |
| S1 | 3 | 3 | 3.5 | 1.75 | 1 | 3 |
| S2 | 4 | 2 | 2 | 2.25 | 2.5 | 3 |
| S3 | 1.5 | 2.5 | 1 | 2 | 2.5 | 1 |
| S4 | 1.5 | 2.5 | 3.5 | 4 | 4 | 3 |

Results in Tables 4 and 5 indicate that the best models in terms of prediction accuracy are **Models NB** and **Pois**, since they had better predictions in the validation set based on both the posterior predictive checks (Cor and MSEP) implemented, although in terms of goodness of fit, **Model LN** was the best. These results are in partial agreement with the findings of Montesinos-Lopez et al. (2015), who came to the conclusion that **Models NB** and **Pois** are good alternatives for modeling count data, although in this study, the best predictions were produced by **Model LN**. However, this model did not take into account the *G × E* interaction.

## Discussion

Developing specific methods for count data for genome-enabled prediction can help to improve the selection of candidate genotypes early in time when the phenotypes are counts. However, currently in genomic selection, phenotypic data (dependent variable) are not taken into account before deciding on the modeling approach to be used, mainly due to the lack of genome-enabled prediction models for non-normal phenotypes. The Bayesian regression models proposed in this paper aim to fill this lack of genome-enabled prediction models for non-normal data.

The first advantage of our proposed methods for count data is that they take into account the nonlinear relationship between responses and consider the specific properties of counts, including discreteness, non-negativity, and over-dispersion (variance greater than the mean); this guarantees that the predictive response will not be negative, which makes no sense for count data. In addition, our methods take into account *G × E*, which plays a central role when selecting candidates genotypes in plant breeding.

Another advantage of our proposed method is that the proposed Gibbs sampler has an analytical solution since we were able to obtain all the full conditional distributions required analytically. This was possible because we constructed our Gibbs sampler using the data augmentation approach proposed by Polson et al., (2013) for count data. For this reason, we believe it is an attractive alternative for fitting complex multilevel data for counts because, in addition to its simplicity, it can generate samples from a high dimensional probability distribution.

Our proposed methods showed superior performance in terms of prediction accuracy compared to **Models Normal** and **Log-Normal**. Also, we observed that in **Models NB** and **Pois** taking into account the *G × E* increase considerable the prediction accuracy which is expected since there is enough scientific evidence that including the *G × E* interaction improve prediction accuracy. Finally, more research is needed to study the proposed methods using real data sets and to extend the proposed genomic-enabled prediction models to deal with so many zeros in count response variables and for modeling multiple traits.

## Acknowledgments

We very much appreciate CIMMYT field collaborator, laboratory assistants, and technicians who collected the phenotypic and genotypic data used in this study.

## Appendix A

Derivation of full conditional distribution for all parameters.

### <u>Full</u> conditional for β^*^

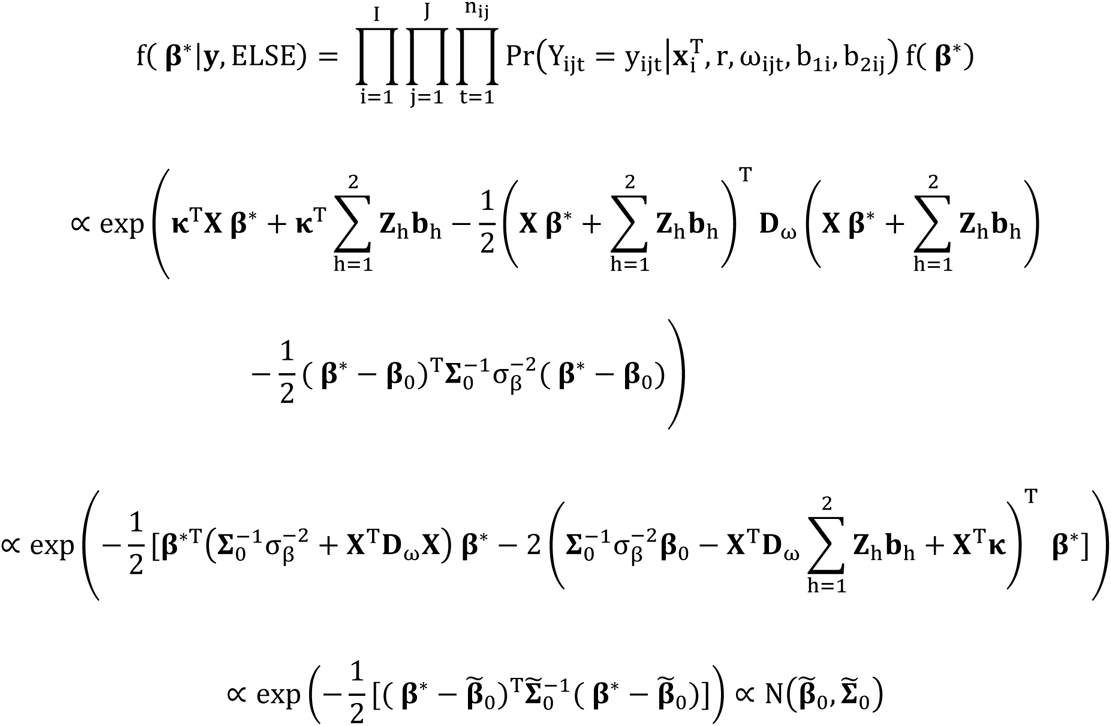

where 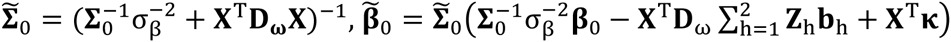.

### <u>Full conditional for</u> ω_ijt_

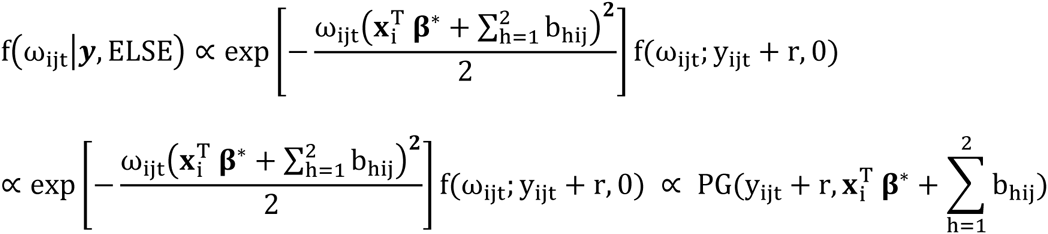

### <u>Full conditional for</u> b_1_

Defining **η**^1^ = **X β**\* + ***Z***_2_**b**_2_ the conditional distribution of **b**_1_ is given as

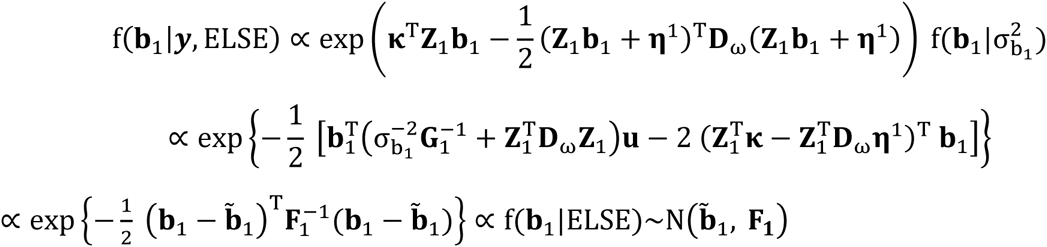

where 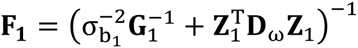 and 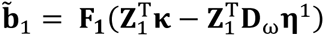.

### <u>Full conditional for</u> 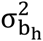

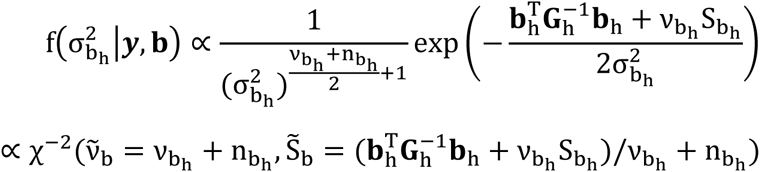

with 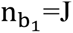 and 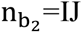.

### <u>Full conditional for</u> 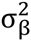

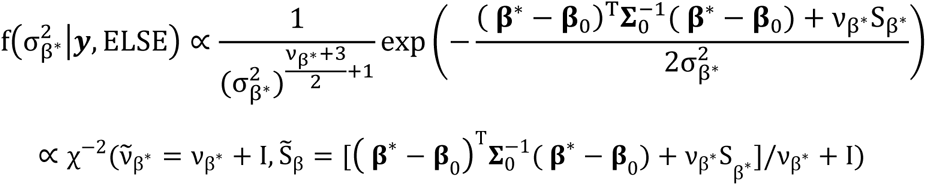

### <u>Full conditional for</u> r

To make the inference of r, we first place a gamma prior on it as r~G(a_0_,1/b_0_). Then we infer a latent count L for each count conditional on Y and r. To derive the full conditional of r, we use the following parameterization of the NB distribution: Y ~ NB(π, r) with 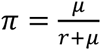. Since L~ Pois(−r log(1 − π)), by construction we can use the Gamma-Poisson conjugacy to update r. Therefore,

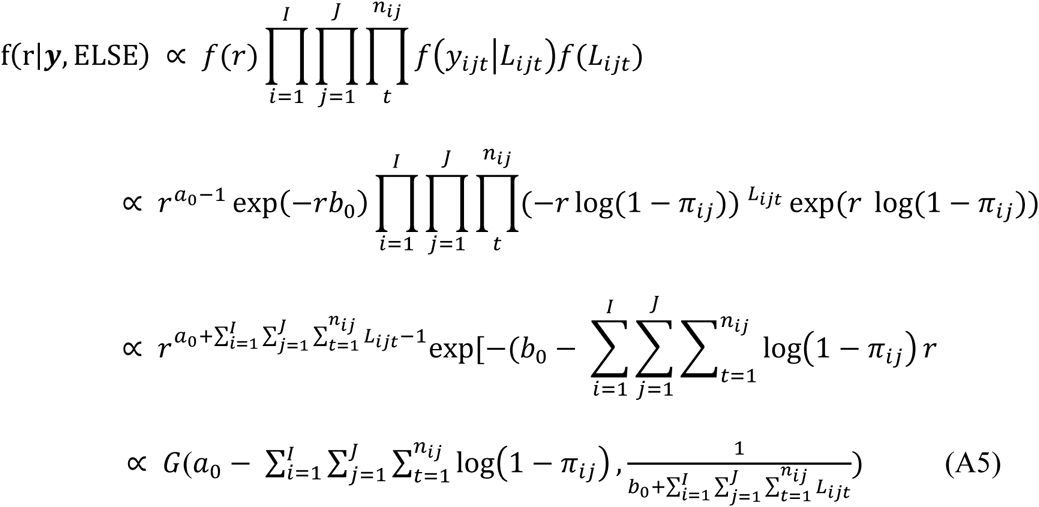

According to Zhou et al. (2012), the conditional posterior distribution of *L_ijt_* is a Chinese restaurant table (CRT) count random variable. That is, *L_ijt_~CRT*(*y_ijt_*,*r*) and we can sample it as 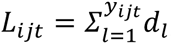, where 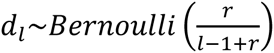. For details of the CRT random variable derivation, see Zhou and Carin (2012, 2015).

## References

Cameron, A.C., and P.K. Trivedi. (1986). Econometric models based on count data. Comparisons and applications of some estimators and tests. Journal of Applied Econometrics 1(1): 29–53.

Cavanagh, C.R., Chao, S., Wang, S. et al. (2013) Genome-wide comparative diversity uncovers multiple targets of selection for improvement in hexaploid wheat landraces and cultivars. Proceedings of the National Academy of Sciences. 110(20) 8057–8062.

de los Campos, G., A.I. Vazquez, R. Fernando, Y.C. Klimentidis, and D. Sorensen. (2013). Prediction of complex human traits using the genomic best linear unbiased predictor. PLoS Genetics 9: e1003608.

de los Campos G, Pataki A, Pérez P (2014). The BGLR (Bayesian Generalized Linear Regression) R-Package. [http://bglr.r-forge.r-project.org/BGLR-tutorial.pdf

Falconi-Castillo, E. (2014). Association mapping for detecting QTLs for Fusarium head blight and Yellow rust resistance in bread wheat. Ph.D. Dissertation. Michigan State University, East Lansing, Michigan, USA.

Garrod, A.E. (1902). The incidence of alkatonuria: a study in chemical individuality. Lancet 160: 1616–1620.

Gelfand, A.E., and A.F. Smith. (1990). Sampling-based approaches to calculating marginal densities. J. Am. Statist. Assoc. 85: 398–409.

Geyer, C.J. (1992). Practical Markov Chain Monte Carlo. Stat Sci 473–483.

Goddard, M.E., and B.J. Hayes. (2009). Mapping genes for complex traits in domestic animals and their use in breeding programmes. Nat. Rev. Genet. 10: 381–391.

Jiao, S, L. Hsu, S. Bézieau, H. Brenner, A.T. Chan, et al. (2013). SBERIA: Set Based Gene-Environment Interaction test for rare and common variants in complex diseases. Genet. Epidemiol. 37: 452–64.

Kraft, P., Y.C. Yen, D.O. Stram, J. Morrison, and W.J. Gauderman. (2007). Exploiting gene environment interaction to detect genetic associations. Hum. Hered. 63: 111–119.

Link, W.A., and M.J. Eaton (2012). On thinning of chains in MCMC. Methods Ecol Evol 3: 112–115.

MacEachern, S.N., and L.M. Berliner. (1994). Subsampling the Gibbs sampler. Am. Stat. 48: 188–190.

Montesinos-López, O.A., A. Montesinos-López, P. Pérez-Rodríguez, G. de los Campos, K.M. Eskridge et al. (2015a). Threshold models for genome-enabled prediction of ordinal categorical traits in plant breeding. G3|Genes|Genomes|Genetics 5: 1–10.

Montesinos-López, O.A., A. Montesinos-López, J. Crossa, J. Burgueño, and K.M. Eskridge. (2015b). Genomic-enabled prediction of ordinal data with Bayesian logistic ordinal regression. G3|Genes|Genomes|Genetics 5: 2113–2126.

Montesinos-López, O.A., A. Montesinos-López, P. Pérez-Rodríguez, K.M. Eskridge, X. He, P. Juliana, P. Singh, and J. Crossa. (2015c). Genomic prediction models for count data. Journal of Agricultural, Biological, and Environmental Statistics (JABES). 20: 533–554, DOI: 10.1007/s13253-015-0223-4.

Murcray, C.E., J.P. Lewinger, and W.J. Gauderman. (2009). Gene-environment interaction in genome-wide association studies. Am. J. Epidemiol. 169: 219–226.

Pérez-de-Castro, A.M., S. Vilanova, J. Cañizares, L. Pascual, J.M. Blanca, M.J. Diez, and B. Picó. (2012). Application of genomic tools in plant breeding. Current Genomics 13(3): 179.

Polson, N.G., J.G. Scott, and J. Windle. (2013). Bayesian inference for logistic models using Pólya-Gamma latent variables. J. Am. Statist. Assoc. 108: 1339–1349.

Quenouille, M.H. (1949). A relation between the logarithmic, Poisson, and negative binomial series. Biometrics 5: 162–164.

R Core Team (2015). R: A language and environment for statistical computing. R Foundation for Statistical Computing. Vienna. Austria. ISBN 3-900051-07-0. URL http://www.R-project.org/.

Stroup, W.W. (2015). Rethinking the analysis of aon-Normal data in alant and soil science. Agron. J. 107: 811–827.

Teerapabolarn, K., and K. Jaioun. (2014). An improved Poisson approximation for the Negative binomial bistribution. Applied Mathematical Sciences 8(89): 4441–4445.

Thomas, D. (2011). Response to ‘Gene-by-environment experiments: a new approach to finding the missing heritability’ by Van Ijzendoorn et al. Nature Rev. Genet. 12: 881.

Turesson, G. (1922). The genotypical response of the plant species to the habitat. Hereditas 3: 211–350.

Van Os, J., and B. Rutten (2009). Gene-environment-wide interaction studies in psychiatry. Am J Psychiatry 166: 964–966.

VanRaden, P.M. (2008). Efficient methods to compute genomic predictions. J. Dairy Sci. 91: 4414–4423.

Windle, J., C.M. Carvalho, J.G. Scott, L. Sun. (2013). Polya-Gamma Data Augmentation for Dynamic Models. arXiv preprint arXiv:1308.0774.

Winham, S.J., and J.M. Biernacka. (2013). Gene-environment interactions in genome-wide association studies: current approaches and new directions. J. Child Psychol. Psychiatry 54: 1120–1134.

Yaacob, W.F.W., M.A. Lazim, and Y.B. Wah. (2010). A practical approach in modelling count data. In Proceedings of the Regional Conference on Statistical Sciences (pp. 176–183).

Zhang, Z., U. Ober, M. Erbe, H. Zhang, N. Gao, et al. (2014). Improving the accuracy of whole genome prediction for complex traits using the results of genome wide association studies. PloS One 9: e93017.

Zhou, M., L. Li, D. Dunson, and L. Carin. (2012). Lognormal and gamma mixed negative binomial regression. In Machine Learning: Proceedings of the International Conference on Machine Learning (vol. 2012. p. 1343). NIH Public Access.

Zhou, M., and L. Carin. (2012). Augment-and-conquer negative binomial processes. In Advances in Neural Information Processing Systems (pp. 2546–2554).

Zhou, M., and L. Carin. (2015). Negative binomial process count and mixture modeling. Pattern Analysis and Machine Intelligence, IEEE Transactions on, 37(2): 307–320.

